# Genetic dissection of the fermentative and respiratory contributions supporting *Vibrio cholerae* hypoxic growth

**DOI:** 10.1101/2020.04.27.065367

**Authors:** Emilio Bueno, Brandon Sit, Matthew K. Waldor, Felipe Cava

## Abstract

Both fermentative and respiratory processes contribute to bacterial metabolic adaptations to low oxygen tension (hypoxia). In the absence of O_2_ as a respiratory electron sink, many bacteria utilize alternative electron acceptors such as nitrate (NO_3_^−^). During canonical NO_3_^−^ respiration, NO_3_^−^ is reduced in a stepwise manner to N_2_ by a dedicated set of reductases. *Vibrio cholerae,* the etiological agent of cholera, only requires a single periplasmic NO_3_^−^ reductase (NapA) to undergo NO_3_^−^ respiration, suggesting that the pathogen possesses a non-canonical NO_3_^−^ respiratory chain. Here, we used complementary transposon-based screens to identify genetic determinants of general hypoxic growth and NO_3_^−^ respiration in *V. cholerae*. We found that while the *V. cholerae* NO_3_^−^ respiratory chain is primarily composed of homologues of established NO_3_^−^ respiratory genes, it also includes components previously unlinked to this process, such as the Na+-NADH dehydrogenase Nqr. The ethanol-generating enzyme AdhE was shown to be the principal fermentative branch required during hypoxic growth in *V. cholerae*. Relative to single *adhE* or *napA* mutant strains, a *V. cholerae* strain lacking both genes exhibited severely impaired hypoxic growth *in vitro* and *in vivo.* Our findings reveal the genetic bases for interactions between disparate energy production pathways that support pathogen fitness in shifting conditions. Such metabolic specializations in *V. cholerae* and other pathogens are potential targets for antimicrobial interventions.

**IMPORTANCE:** Bacteria reprogram their metabolism in environments with low oxygen levels (hypoxia). Typically, this occurs via regulation of two major, but largely independent, metabolic pathways-fermentation and respiration. Here, we found that the diarrheal pathogen *Vibrio cholerae* has a respiratory chain for NO_3_^−^ that consists largely of components found in other NO_3_^−^ respiratory systems, but also contains several proteins not previously linked to this process. Both AdhE-dependent fermentation and NO_3_^−^ respiration were required for efficient pathogen growth in both laboratory conditions and in an animal infection model. These observations provide genetic evidence for fermentative-respiratory interactions and identify metabolic vulnerabilities that may be targetable for new antimicrobial agents in *V. cholerae* and related pathogens.

---

Respiration generates most of the energy produced in the cell, making it a vital process to most organisms. Although the general mechanism of respiration is well-conserved, many bacteria employ specialized respiratory machineries for adaptation to diverse environmental conditions. One such condition that bacteria frequently encounter is the lack of oxygen, also known as hypoxia/anoxia. When oxygen is present, most bacteria use molecular O_2_ as the terminal electron acceptor to drive aerobic respiration and complete the electron transport chain (ETC). When O_2_ levels are insufficient to support respiration, some bacteria and archaea can still grow by using certain alternative electron acceptors (AEAs) in the place of O_2_ (1, 2). AEAs, which include nitrogen oxides (e.g. nitrate (NO_3_^−^)), dimethyl sulphoxide (DMSO), trimethylamine N-oxide (TMAO), and fumarate, are used to terminate anaerobic ETCs. Like their aerobic counterparts, anaerobic ETCs are initiated with electrons donated by membrane-bound NADH dehydrogenases and consist of a series of redox reactions involving quinones and other electron carrier proteins. Anaerobic ETCs are highly specialized, as specific conditions may preferentially employ certain NADH dehydrogenases over others, and each AEA has its own specific associated reductase enzyme or enzyme complex. For example, in *E. coli*, the electrogenic NADH dehydrogenase I (NDH-1) is the respiratory complex utilized in the presence of fumarate or DMSO. However, when the electron acceptor is NO_3_^−^ or O_2_, the ETC is mainly driven by the non-electrogenic NADH dehydrogenase II (NDH-2) (1, 2).

Anaerobic respiration of NO_3_^−^ generally results in the production of N_2_ gas or NH_4_^+^. NO_3_^−^ reduction to N_2_ (i.e. denitrification) occurs in four stages where NO_3_^−^ is reduced stepwise to N_2_ (NO_3_→NO_2_→NO→N_2_O→N_2_) (3) by nitrate-, nitrite-, nitric oxide-, and nitrous oxide reductases, encoded by the *nar/nap*, *nir*, *nor*, and *nos* genes, respectively (reviewed in (4, 5)). In addition to the importance of NO_3_^−^ respiration in bacteria under free-living conditions, recent studies have also illuminated the role of nitrate respiration during colonization of host tissues, such as the intestines, which lack oxygen (6–8).

When both O_2_ and an AEA are absent, bacteria are still able to generate energy by undergoing fermentation, where energy is produced by substrate-level phosphorylation, and pyridine nucleotides replace respiratory quinones as intermediate electron drivers. Fermentation is a bipartite process where glucose oxidation generates NADH and pyruvate, which is subsequently reduced to regenerate NAD^+^. The NAD^+^ regeneration step is comprised of multiple branches with dedicated enzymes, which produce small organic byproducts such as acetate, formate, lactate and ethanol (8–10). Specifically, reduction of pyruvate to lactate or ethanol provides a mechanism for NAD^+^ regeneration, while the acetate branch generates energy in the form of ATP (1, 2). Similarly to anaerobic respiration, fermentation is known to be critical for pathogens to colonize oxygen-scarce host niches (11, 12).

*Vibrio cholerae*, the causative agent of the diarrheal disease cholera, is a facultative anaerobic bacterium (13) that can be found in marine environments or associated with human hosts. In humans, *V. cholerae* colonizes the small intestine, where production of cholera toxin induces secretory diarrhoea, expelling the pathogen to the environment, where it can be ingested by a new host to complete a transmission cycle (14, 15). Transition from the human gastrointestinal tract to the aquatic environment and vice versa requires the pathogen to adapt to rapidly fluctuating oxygen levels. As part of this adaptation, in addition to an intact fermentation machinery (16), *V. cholerae* encodes three terminal oxidases with different affinities for oxygen to fine-tune aerobic respiration (16) and three AEA reductases to allow it to anaerobically respire fumarate, TMAO and NO_3_^−^ (17, 18). *V. cholerae* also encodes a variety of dehydrogenases that may function in respiratory metabolism, but their exact roles and whether they have functions specialized to different AEAs remains unknown.

In contrast to other NO_3_^−^-respiring bacteria, *V. cholerae* only encodes the periplasmic nitrate reductase Nap (*napFDABC,* VC0676-VC0680) and does not harbor any of the downstream reductases (16). As NO_2_^−^ accumulation is growth-inhibitory, this would seemingly restrict its ability to use NO_3_^−^ as an AEA. However, we recently reported that *V. cholerae* can in fact undergo NO_3_^−^ respiration, and that incomplete NO_3_^−^ reduction allows the bacterium to control its fitness in hypoxic environments in the face of differing pH levels (8). Here, we decipher the complete set of *V. cholerae* loci required for its hypoxic metabolism and describe how fermentation and NO_3_^−^ respiration interact to support pathogen fitness in the laboratory and during infection.

## RESULTS

### Genetic screens for determinants of hypoxic fitness and NO_3_^−^ respiration in *V. cholerae*

To uncover genetic determinants of NO_3_^−^ respiration in *V. cholerae,* we performed transposon-insertion sequencing (TIS), a high-throughput technique for identifying the loci required for bacterial fitness in a given selective condition. We first attempted to compare pooled transposon mutant libraries of *V. cholerae* C6706 in hypoxic respiratory (+NO_3_^−^, −O_2_, pH 8) and non-respiratory (−NO_3_^−^, −O_2_, pH 8) conditions, but encountered technical limitations that precluded high-resolution analysis of these screens. Instead, we compared libraries grown in hypoxic respiratory (+NO_3_^−^, −O_2_, pH 8) or typical laboratory conditions (−NO_3_^−^, +O_2_, pH 8), enabling identification of the *V. cholerae* gene set required for hypoxic survival in the presence of an AEA (NO_3_^−^) (Fig. 1A). A relatively large number of genes (256) were identified as under-represented by this screen, indicating *V. cholerae* requires an extensive gene set to adapt to O_2_-scarcity in these conditions. Strikingly, many under-represented loci identified in this screen were genes that are known to function in NO_3_^−^ respiration in other bacteria, such as *napA, fnr, narP and moaA*, supporting the idea that NO_3_^−^ respiration was a strong selective force in the screen.

**Figure 1.**
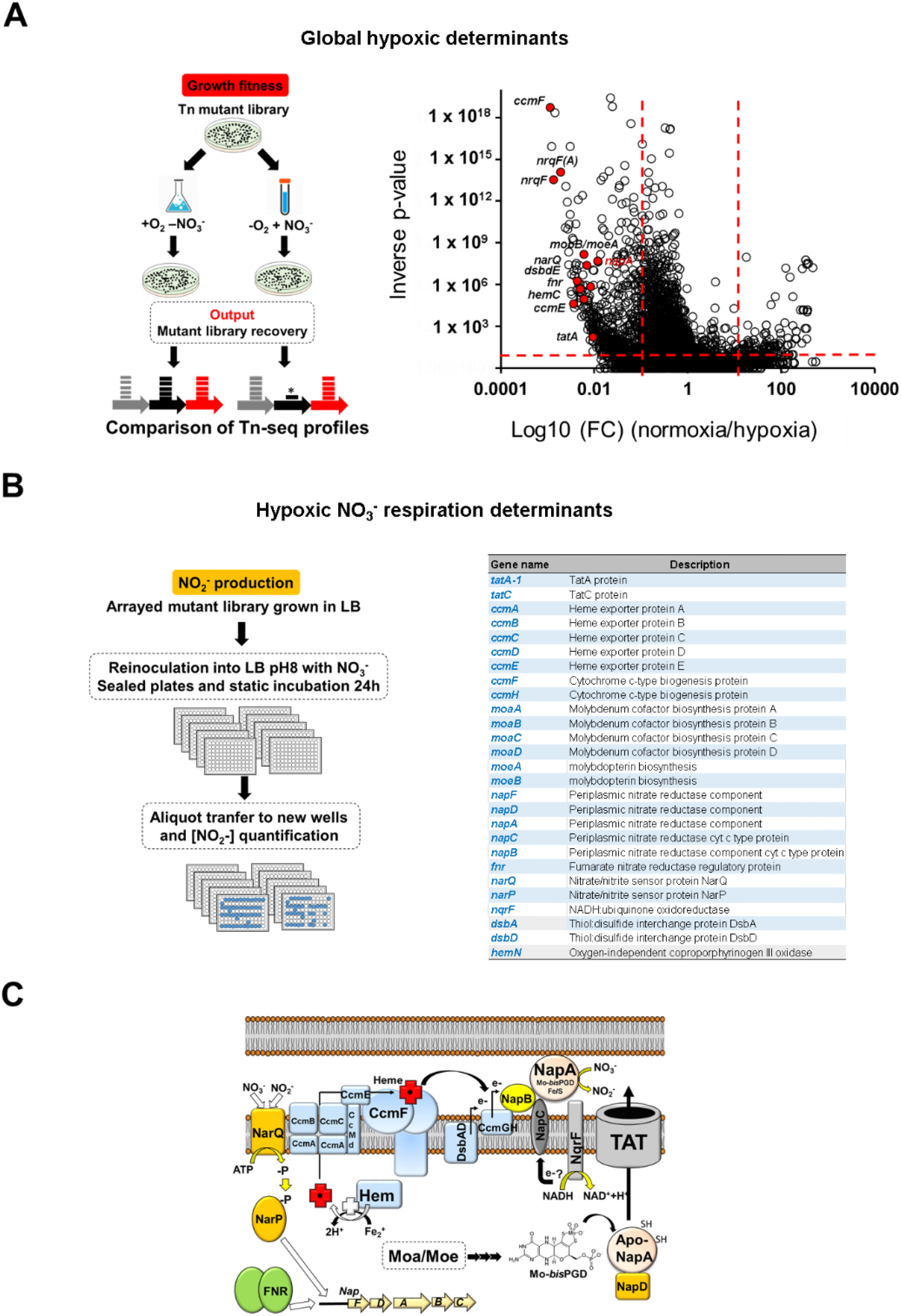
Identification of *V. cholerae* genetic determinants for hypoxic metabolism and NO_3_^−^ reduction. (A) Left: Experimental workflow for TIS screen comparing *V. cholerae* mutant libraries grown in normoxic or hypoxic conditions. Right: volcano plot of Con-ARTIST analysis output depicting read fold change (FC) and inverse Mann-Whitney U p-value for each gene queried in the TIS screen. Red dotted lines indicate arbitrary thresholds of FC >10 or <0.1 and an inverse p-value >20. Solid red circles indicate genes known to be involved in NO_3_^−^ respiration. (B) Left: Experimental workflow for arrayed transposon mutant screen to identify genes required for NO_3_^−^ reduction to NO_2_^−^ in *V. cholerae.* Right: list of mutant strains queried in the arrayed screen (see Supplementary table 1 for an extended version. (C) Proposed organization of the NO_3_^−^ reduction machinery in *V. cholerae* based on results from the transposon screens.

To determine from the pooled TIS screen exactly which genes were required for general hypoxic fitness (e.g., genes required for oxygen sensing or fermentation) or specifically required for NO_3_^−^ respiration, we next performed a complementary arrayed transposon screen with a non-redundant C6706 insertion mutant library (19) (Fig. 1B, left). In contrast with the pooled screen, where the readout was the relative fitness of mutants in each condition, this screen directly evaluated the capacity of individual mutants to produce nitrite (NO_2_^−^), the result of NO_3_^−^ respiration. From the arrayed screen, 24 candidates were implicated in NO_3_^−^ reduction in *V. cholerae* (Fig. 1B, right), including the *nap* operon, the oxygen response regulatory protein Fnr, the two component system NarQ-NarP, the genes responsible for the synthesis of molybdenum cofactor, the *ccm* and *dsbD* loci implicated in the maturation of *c*-type cytochromes, the coproporphyrinogen III oxidase HemN implicated in the biosynthesis of haem groups, the Tat transport system, and the *V. cholerae* NADH dehydrogenase Na^+^-Nqr. All 24 of these candidates were under-represented in the initial TIS screen, supporting our previous observations that NO_3_^−^ respiration, when active, is critical for *V. cholerae* growth under hypoxia (8).

We previously showed that hypoxic NO_3_^−^ reduction in *V. cholerae* exerts divergent pH-dependent functions (8). In alkaline environments (pH = 8), NO_3_^−^ reduction is coupled to ATP production to support respiratory growth. However, in acidic (pH < 6) conditions, respiratory activity is turned off and instead, NO_3_^−^ reduction slows the negative effects of increased fermentative activity, namely the acute acidification of the extracellular environment and associated loss of culture viability. We hypothesized that mutants in the genes identified by our screens would also show similar pH-dependent phenotypes. Indeed, all tested mutant strains (*napC, tatA, ccmF, narP* and *moaA*) were unable to arrest growth in acidic conditions, leading to excessive reduction of medium pH and total loss of viability (Fig. 2B). Under alkaline conditions (Fig. 2B), the same mutant strains also did not benefit from NO_3_^−^ addition, phenocopying the behaviour of the known NO_3_^−^-reduction and respiration-deficient Δ*napA* strain (8). As all of these genes are known to participate in NO_3_^−^ reduction and respiration in other bacteria, these data suggest that *V. cholerae* uses a largely canonical NO_3_^−^ reduction pathway to control the pH-dependent fitness outcomes of hypoxia.

**Figure 2.**
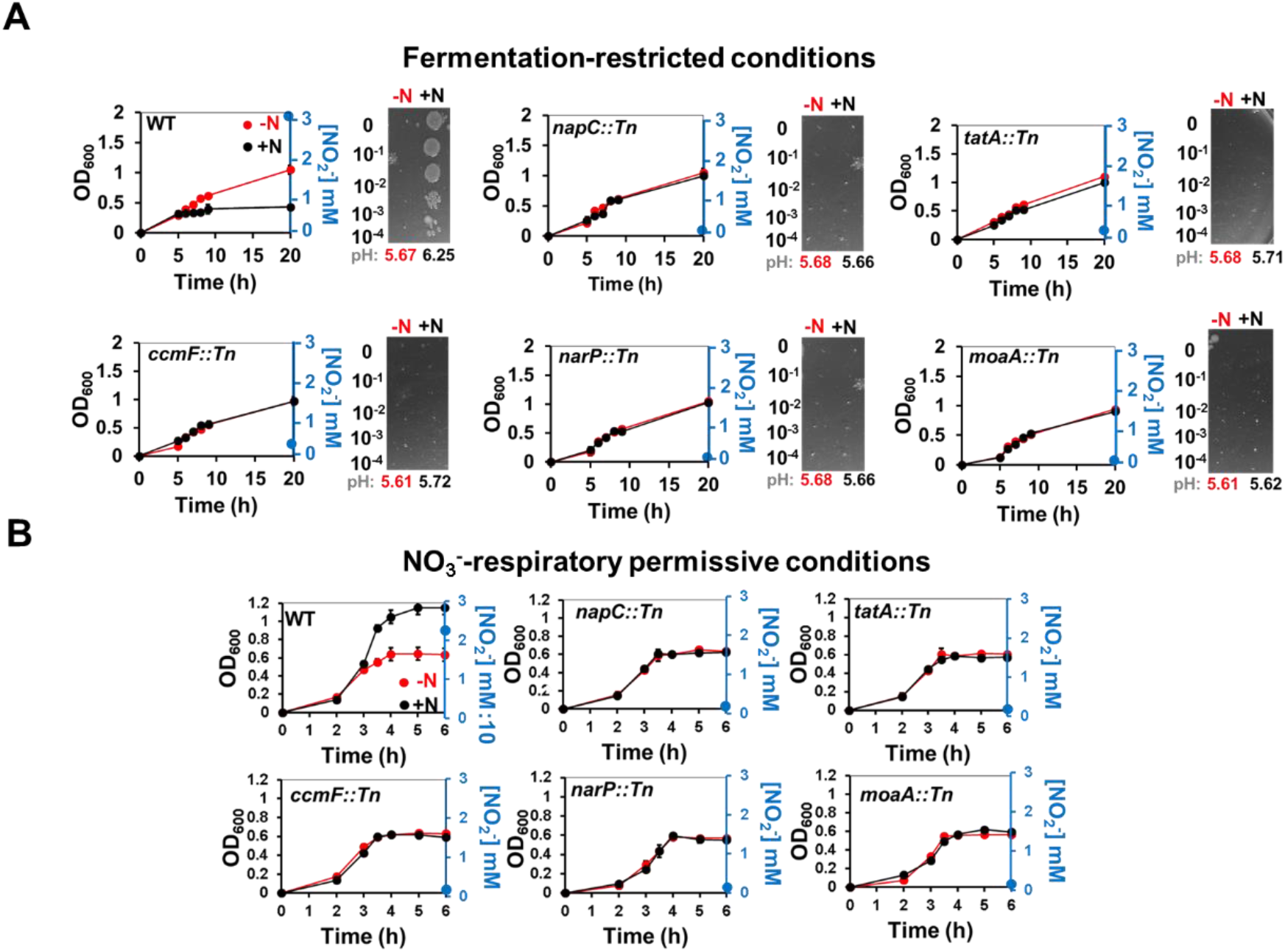
Genes involved in NO_3_^−^ reduction in *V. cholerae* are required for divergent pH-dependent responses to NO_3_^−^ under hypoxia. (A) Optical density (OD_600_) growth curves and endpoint CFU plating for C6706 WT *V. cholerae* and *napC, tatA, ccmF, narP* and *moaA* transposon insertion strains. Cultures were grown hypoxically in M9 minimal media + 1% glucose (fermentative conditions) in the absence (red, −N) or presence (black, +N) of 3 mM NO_3_^−^. Endpoint pH and NO_2_^−^ levels were measured at the time of plating (20 hours). (B) OD_600_ growth curves for strains in Panel A grown in LB at pH 8 (NO_3_^−^ respiratory conditions) with or without 3mM NO_3_^−^. Endpoint NO_2_^−^ accumulation is shown in blue on the right y-axis. Data are the mean of 3 biological replicates +/− SE.

### The Dsb pathway and Na^+^-Nqr dehydrogenase are required for NO_3_^−^ respiration in *V. cholerae*

In addition to the canonical ‘core’ NO_3_^−^ respiratory machinery described above, we identified two other systems in our screen that showed impaired, but not abrogated NO_3_^−^ respiratory growth when disrupted: the Dsb disulfide bond forming system and the Na^+^-dependent NADH dehydrogenase Nqr. Both systems, represented by their essential components *dsbA* and *nqrF*, respectively, have not been specifically implicated in NO_3_^−^ respiration. To query these subtler phenotypes, we switched from tube cultures to a plate-reader based growth assay and used mineral oil overlays combined with oxygen-impermeant plate sealing film to establish stable anoxic conditions. In these conditions, we still observed that WT *V. cholerae* experienced a rapid increase on growth in pH 8 medium when supplemented with NO_3_^−^, confirming that NO_3_^−^ respiration is induced in this system (Fig. 3A). The magnitude of this phenotype was significantly reduced in the *dsbA*-defective strain (Fig. S1). Since the Dsb pathway maintains redox homeostasis, we attempted to alter the redox state of WT *V. cholerae* by adding glutathione to the cultures. Addition of glutathione phenocopied the attenuated NO_3_^−^ respiration of Δ*dsbA* (Fig. S1), suggesting that in addition to hypoxia and the presence of NO_3_^−^, cellular redox status is another important factor controlling the induction of NO_3_^−^ reduction (and respiration) in *V. cholerae*.

**Figure 3.**
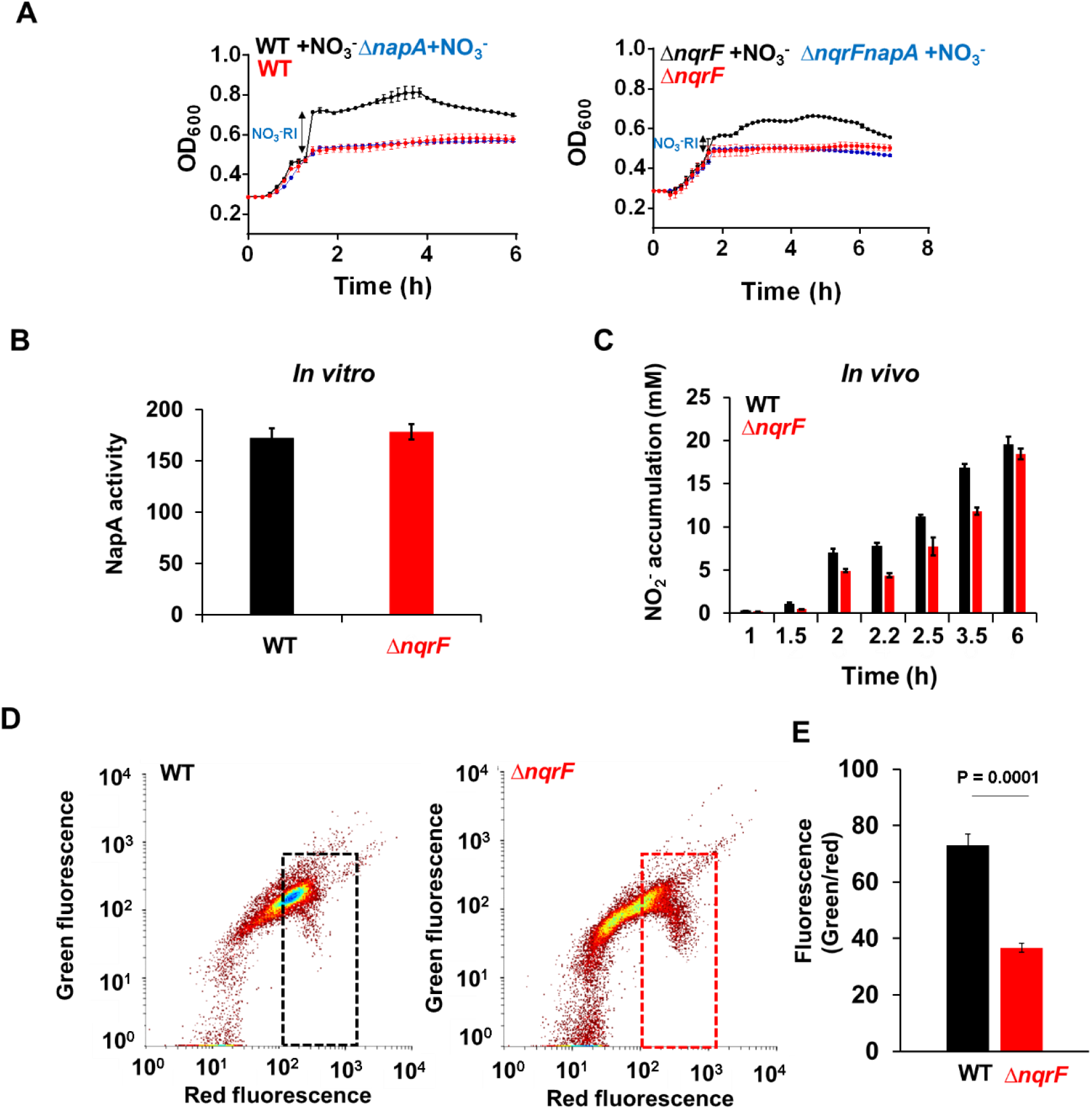
The NADH dehydrogenase Na^+^-Nqr participates in NO_3_^−^ respiration in *V. cholerae*. (A) OD_600_ growth curve of *V. cholerae* WT, Δ*napA, ΔnqrF* and Δ*nqrF/napA* strains grown hypoxically in LB medium buffered at pH8 in the presence (black and blue) or absence (red) of 20 mM NO_3_^−^. Induction of the nitrate respiratory pathway is labeled as NO_3_^−^-RI in each growth curve. (B) Methyl viologen-dependent *in vitro* nitrate reductase (NR) activity (nmol NO_2_^−^ produced/min/mg protein). Aliquots from cultures in Panel A were collected after 2 hours of incubation and assayed for NR activity. (C) Time-course *in vivo* NR activity. (D) Representative membrane potential flow cytometry plots from DiOC2-stained cells. (E) Quantification of green/red fluorescence ratios in cells indicated by dashed boxes in Panel D. Data are the mean of 3 biological replicates +/− SE. Statistical significance was determined by the Student’s t-test.

Akin to Δ*dsbA*, the growth kinetics of Δ*nqrF V. cholerae* in the presence or absence of NO_3_^−^ under hypoxia indicated that NO_3_^−^ respiratory chain activity was approximately half that of the WT (Fig. 3A). To understand the mechanism of this defect, we asked whether the activity of NapA, the terminal NO_3_^−^ reductase, was altered in Δ*nqrF* cells. Although NapA activity *in vitro* was unaltered (Fig. 3B panel), *in vivo* NapA activity, measured as NO_2_^−^ release into the culture supernatant, was decreased ~2x relative to wild type (Fig. 3B right panel). Na^+^-Nqr is thought to initiate the ETC by coupling dehydrogenase activity to proton export to the periplasm, resulting in electrons shunted to the ubiquinone pool with a concomitant maintenance of membrane potential. To determine if Na^+^-Nqr maintains membrane potential during NO_3_^−^ respiration, we used the indicator dye DiOC_2_(3) (20), which diffuses through bacterial membranes and accumulates within the cytoplasm depending on the cellular membrane potential. When DiOC_2_(3) is transferred into the cytoplasm at low concentrations (due to low membrane potential) it exhibits green emission. However, when membrane potential is high, DiOC_2_(3) concentrates and self-associates, shifting to red emission. We found that membrane potential, while intact, was reduced by about half in the Δ*nqrF* strain (Fig. 3D), a magnitude consistent with the decreased levels of NO_3_^−^ reduction in this mutant and suggestive of an electrogenic electron donor role for Na^+^-Nqr in NO_3_^−^ respiration.

Although inactivation of Na^+^-Nqr markedly attenuated the electrons transferred into the Nap system, the incomplete inhibition of NO_3_^−^ reduction and growth under NO_3_^−^ reducing conditions suggests the existence of other electron-donating factors. To address this, we took a candidate-based approach and inactivated four putative NADH dehydrogenases: the NADH-2 dehydrogenase (VC1890), d-amino acid dehydrogenase (VC0786), putative dehydrogenase VC1581, and the NO_3_^−^-inducible formate dehydrogenase FdnI (VC1511) (21) (Fig. 4A). All dehydrogenase mutant strains were able to grow by NO_3_^−^ respiration (Fig. 4B, left column) to a similar level as WT *V. cholerae* (Fig. 3A, left panel). To test whether any of these dehydrogenases could be working in concert with NqrF, we constructed double mutants in the Δ*nqrF* background. Strikingly, NO_3_^−^ respiration in all of the double mutants (Fig. 4, right column) still phenocopied the *nqrF* single deletion strain (Fig 3A), indicating that Na^+^-Nqr is the primary dehydrogenase that contributes electrons into the *V. cholerae* NO_3_^−^ respiration system.

**Figure 4.**
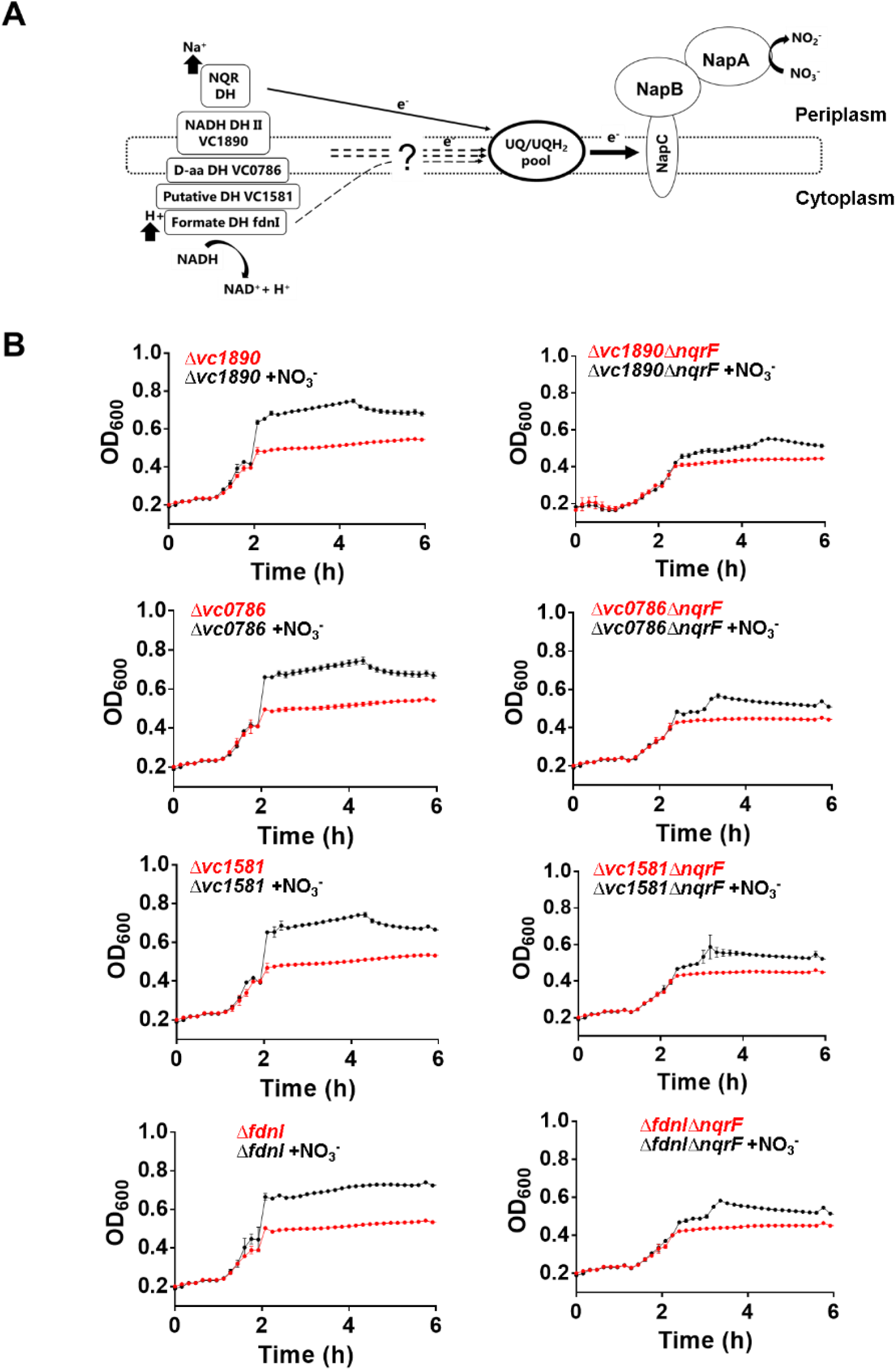
VC1890, VC0786, VC1581 and VC1511 do not contribute to NO_3_^−^ respiration in *V. cholerae*. (A) Proposed schematic of NO_3_^−^-terminated electron transport chain in *V. cholerae* in hypoxic NO_3_-replete conditions. Redox equivalents are channeled into the ETC by one or more membrane-bound dehydrogenases that feed the ubiquinone (UQ) pool. Electrons from the UQ pool are ultimately transferred to NO_3_^−^ by specific terminal reductases. (B) OD_600_ growth curves of *V. cholerae* WT and dehydrogenase-deficient strains. Each dehydrogenase was also deleted in combination with Na^+^-Nqr (right panels). Strains were grown hypoxically in LB medium buffered at pH8 in the presence (black) or absence (red) of 20 mM NO_3_^−^. Data are the mean of 3 biological replicates +/− SE.

### Interactions between NO_3_^−^ respiration and AdhE-driven fermentation support hypoxic fitness in*V. cholerae*

Unexpectedly, our TIS screen did not identify any genes known to be involved in fermentation, a metabolic process that we expected would be important for growth under hypoxia (Fig. 1, 5A). We hypothesized this was due to metabolic redundancy between respiration and fermentation, obscuring the contribution of the latter to hypoxic fitness when an AEA such as NO_3_^−^ is present. To understand this phenomenon, we inactivated fermentation by deleting the genes encoding lactate dehydrogenase LdhA, acetate kinase Ack or alcohol dehydrogenase AdhE (Fig. 5A). While hypoxic growth with or without NO_3_^−^ in *ack* and *ldhA* mutant strains was comparable to WT *V. cholerae*, inactivation of *adhE* resulted in reduced fitness during fermentation-restricted conditions (Fig. 5B). Interestingly, addition of NO_3_^−^ to Δ*adhE* completely restored its growth to WT levels, indicating that during hypoxic nitrate respiratory conditions, fermentative activity is not essential for *V. cholerae* growth (Fig. 5B). We tested whether such compensation of fermentation by respiration is applicable to other bacteria by repeating similar analyses in *Escherichia coli.* Similar to *V. cholerae,* growth impairment of a Δ*adh E. coli* strain in fermentation-restricted conditions was complemented when NO_3_^−^ was added to the medium (Fig. 5C). Thus crosstalk between NO_3_^−^ respiration and fermentation under hypoxia is not restricted to *V. cholerae*.

**Figure 5.**
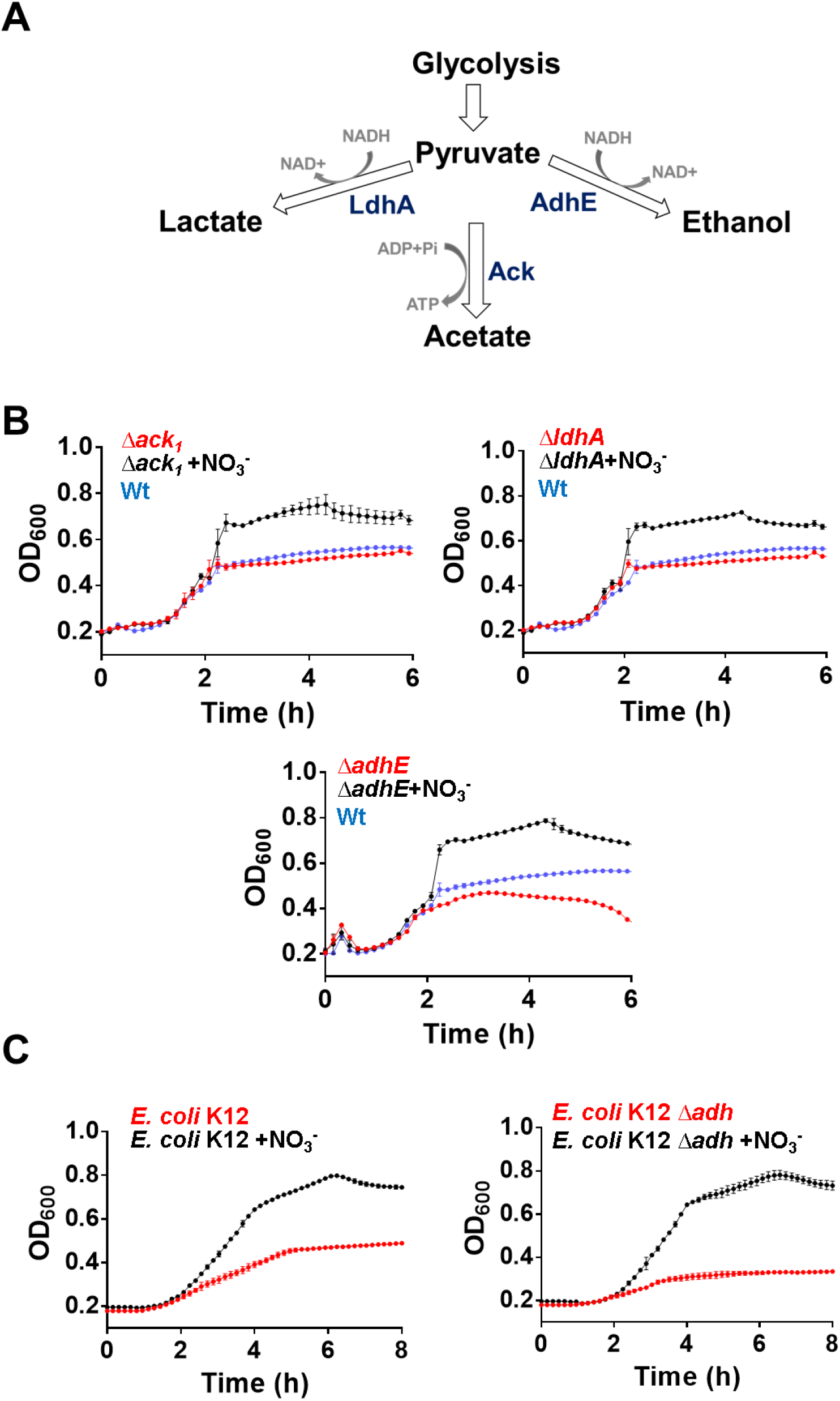
*V. cholerae* fermentation is driven by AdhE. (A) Schematic of fermentative branches in *V. cholerae*. (B) OD_600_ growth curve of *V. cholerae* WT and fermentative mutant strains: Δ*ack1(left), ΔldhA (right), and ΔadhE (bottom)*. (C) OD_600_ growth curve of *E. coli* K12 WT and Δ*adh* strains. Cultures were grown hypoxically in LB medium buffered at pH8 in the presence (black) or absence (red and blue) of 20 mM NO_3_^−^. Data are the mean of 3 biological replicates +/− SE.

To test if similar interactions between fermentation and NO_3_^−^ respiration occurred during competitive growth conditions, we co-cultured WT and mutant *V. cholerae* strains in the presence and absence of NO_3_^−^. In the absence of NO_3_^−^, Δ*adhE* was unable compete efficiently with the WT strain (Fig. 6A). Addition of NO_3_^−^ significantly ameliorated this competitive defect, similarly to the results from single strain growth experiments (Fig. 5B, 6A). When NO_3_^−^ was present, inactivation of both *adhE* and *napA* resulted in a greater loss of competitive fitness vs the WT strain than either single mutant alone, suggesting that both NO_3_^−^ respiration and fermentation support fitness under these conditions. Finally, we performed similar competition studies in the streptomycin-treated adult mouse model of *V. cholerae* intestinal colonization (22). In this model, streptomycin pre-treatment disrupts the normal gut microbiota and raises NO_3_^−^ levels in the intestinal lumen, providing a substrate for NO_3_^−^ respiration-competent bacteria. In previous studies, we found that Δ*napA V. cholerae* had a small but significant (CI ~0.5) defect during competitive infections (8) (reproduced in Fig. 6B). In contrast, the Δ*adhE* mutant did not have a detectable colonization defect, likely because NO_3_^−^ respiration via NapA fully compensated for its lack of fermentative capacity (Fig. 6B). Supporting this hypothesis, we observed that inhibition of the NO_3_^−^ respiratory pathway in Δ*adhE* by inactivation of *napA* resulted in a dramatic loss of competitive fitness against the WT strain (Fig. 6B). Altogether, these results suggest that metabolic interactions between fermentation and respiration mediated by *adhE* and *napA* supports competitive hypoxic fitness in *V. cholerae*.

**Figure 6.**
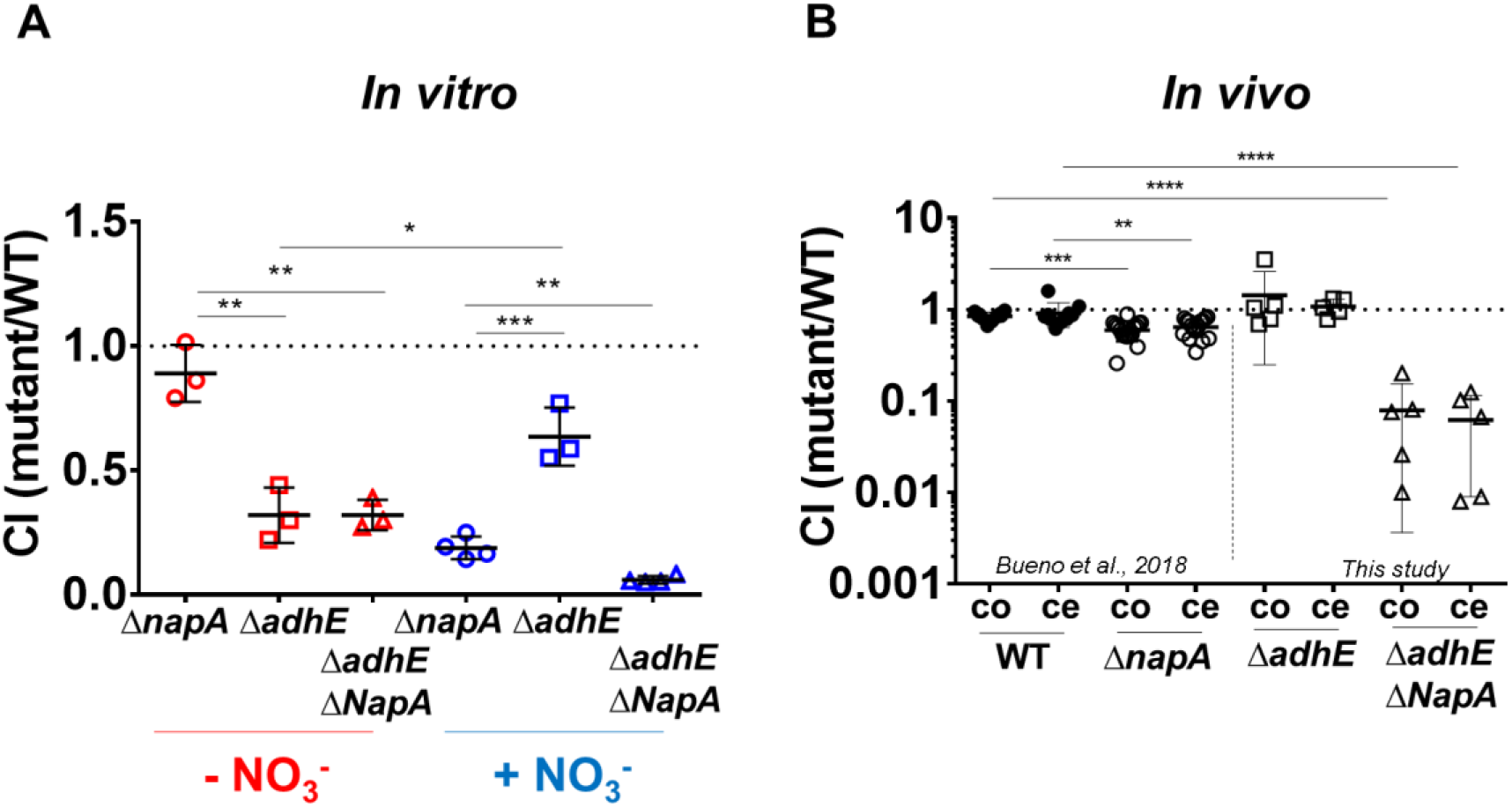
NO_3_^−^ respiration in *V. cholerae* supports lack of fermentation *in vitro* and *in vivo* fitness. (A) *In vitro* competitive indices (CI) of WT *V. cholerae* versus Δ*napA, ΔadhE* or Δ*adhE/napA* strains during hypoxic growth in LB medium buffered at pH 8. CIs were determined at 8 hours post-inoculation. (B) *In vivo* competition of WT *V. cholerae* versus Δ*napA, ΔadhE* and Δ*adhEΔnapA* in streptomycin-treated adult mouse colonization model. The WT control (WT vs. WT *lacZ^−^*) and Δ*napA* competition data (left datasets) are reproduced from reference 8. CIs were determined at 24 hours post-inoculation from harvested colon (co) and cecum (ce) samples. Statistical significance was determined by the Student’s t-test (A) or Mann Whitney-U test (B). A p-value less than 0.05 was considered statistically significant. Bars represent geometric mean + SE.

## DISCUSSION

Most enteric bacterial pathogens encode both NO_3_^−^ respiratory and fermentative enzymes (23–29). However, knowledge regarding the complete composition of these pathways during hypoxic growth is limited. Here, we identified the genetic determinants of NO_3_^−^ respiration and fermentation in the cholera pathogen, *V. cholerae*. The complementary transposon-based approaches, particularly the arrayed mutant screen, allowed us to identify the nitrate respiratory machinery in *V. cholerae.* This system primarily relies on structural and regulatory proteins identified in the NO_3_ ^−^ respiratory systems of other organisms (Fig. 1B). Based on these data, we propose the following scheme for the NO_3_^−^ reduction machinery in *V. cholerae.* First, NO_3_^−^/NO_2_^−^ is sensed by the two-component system NarQP, which cooperates with the O_2_-sensing transcriptional regulator Fnr to activate *napFABCD* operon transcription. To initiate assembly of the mature Nap reductase, newly synthesized NapA is folded by the NapD chaperone and then loaded with the molybdenum cofactor Mo-bisPGD (synthesized by the Moa complex). Holo-NapA is subsequently translocated to the periplasm via the Tat secretion system. Heme moieties for NapB and NapC, which are both cytochrome *c*-type proteins, are likely synthesized by the Hem complex in the cytoplasm. Final cytochrome assembly is performed by Ccm-DsbD in the periplasm, a process that involves thioreduction of apo-NapBC and subsequent covalent ligation of heme to the protein backbone. Once the NapABC complex has matured in the periplasm, NO_3_^−^ can be reduced to NO_2_^−^ by the catalytic subunit NapA.

In addition to this core NO_3_^−^ reduction machinery, we identified two proteins that have not previously been implicated in NO_3_^−^ respiration, including NqrF, a critical subunit of the Na+-translocating NADH:quinone dehydrogenase Nqr. *In V. cholerae*, Na^+^-Nqr has been reported to regulate bacterial motility, metabolism (30), resistance against heavy metals (31), and virulence (32–34). Deletion of Na^+^-Nqr in *V. cholerae* impaired membrane potential and delayed NO_3_^−^ reduction to NO_2_^−^ relative to the WT strain, suggesting that Na^+^-Nqr is required for maintenance of the ETC under NO_3_^−^ respiratory conditions. Inefficient electron transport likely explains the comparatively minor NO_3_^−^-dependent boost in growth of the *nqrF* mutant. We propose that Na^+^-Nqr acts as an initial source of electrons for eventual NO_3_^−^ reduction by Nap, further highlighting the importance of this dehydrogenase in *V. cholerae* biology. Genetic screens in other species, particularly screens of arrayed mutant libraries, should be worthwhile for uncovering additional non-canonical respiratory chain components for NO_3_^−^ reduction and should shed light on the evolution of metabolic pathways.

Although we endeavoured to identify other dehydrogenases that could supplement Na^+^-Nqr-mediated contributions to the NO_3_^−^ respiratory ETC, none of the four candidates tested (VC1890, VC0786, VC1581 and VC1511) exhibited impaired NO_3_^−^ respiratory function, suggesting that there are unannotated *V. cholerae* respiratory dehydrogenases that participate in NO_3_^−^ respiration. We were particularly surprised that VC1890 does not contribute to NO_3_^−^ respiration, as its homologue in *E. coli*, NDH-2, is the main NO_3_^−^-induced NADH dehydrogenase in that species (1, 2). In *V. cholerae,* it seems likely that the activity of several of dehydrogenases, including Na^+^-Nqr, are redundant; thus, inactivation of Na^+^-Nqr together with a single secondary dehydrogenase could be overcome by directing electron flow through additional dehydrogenases. Higher order synthetic genetic analysis in the Δ*nqrF* mutant background combined with transcriptional profiling to find NO_3_^−^-regulated genes that may have dehydrogenase activity will be useful for identification of additional candidates for this function.

An interesting feature of Na^+^-Nqr relative to other electrogenic dehydrogenases is that it exports Na^+^ instead of H^+^, creating a sodium motive force (SMF) rather than a proton motive force (PMF). In the human small intestine, *V. cholerae* secretion of cholera toxin activates intestinal epithelial Cl^−^ ion transporters such as CFTR, leading to massive export of Cl^−^ ions into the intestinal lumen. This is accompanied by concomitant Na^+^ and H_2_O efflux (35). In the context of high concentrations of Na^+^, hypoxia and the presence of NO_3_^−^ within the intestines, Na^+^-Nqr is thus well suited to support *V. cholerae* proliferation and potentially allow the pathogen to compete with commensal bacteria that may lack such Na^+^-dependent machinery. It will be of interest to determine whether Na+-Nqr’s diverse contributions to *V. cholerae* fitness (i.e. metabolism, virulence, and motility) can be untangled experimentally.

Finally, our data have implications for understanding how *V. cholerae* coordinates different metabolic pathways. In general, there is a hierarchical organization of hypoxic bacterial metabolism, where more energetically efficient AEAs such as NO_3_^−^ and TMAO are used first, followed by those that yield less energy (nitrite, DMSO, tetrathionate). Fermentation, the pathway that produces the least energy, is only used if all potential AEAs are absent or have been exhausted from the environment (1, 2). However, in *V. cholerae*, nitrate respiration and fermentation overlap and co-occur, obscuring their individual contributions to growth (8). We found that only the ethanol-generating AdhE-dependent fermentative branch was required during *V. cholerae* fermentative growth under laboratory conditions. Defects in fermentation in Δ*adhE* were complemented in the presence of NO_3_^−^ in a NapA-dependent manner, but a double fermentative/respiratory mutant Δ*adhE*/*napA* exhibited a greater defect than either single mutant, suggesting that *V. cholerae* maintains baseline fermentative activity during NO_3_^−^ respiration. The greater magnitude of the Δ*napA* defect in the presence of NO_3_^−^ than that of Δ*adhE*, suggests that NO_3_^−^ respiration contributes more to fitness in those conditions than fermentation and is the dominant energy production pathway in the presence of NO_3_^−^. This primacy of NO_3_^−^ respiration was corroborated by our *in vivo* studies, where Δ*adhE* displayed no detectable colonization defect, but concomitant inactivation of *napA* and *adhE* lowered the competitive index by more than one order of magnitude. We speculate that *V. cholerae* differentiates between NO_3_^−^-respiratory and non-respiratory conditions by adjusting levels of both its respiratory and fermentative activity in an opposing but non-mutually exclusive manner. Consequentially, NO_3_^−^ respiration in this pathogen coexists with, and energetically complements, fermentation. It will be of interest to determine whether similar relationships exist between fermentation and respiration of non-NO_3_^−^ AEAs, and how the simultaneous presence of more than one suitable AEA will influence those interactions. If these metabolic interactions are conserved in other pathogens besides *V. cholerae,* we propose that respiratory/fermentative crosstalk represents a new class of vulnerability that may be an attractive target for antimicrobial interventions.

## MATERIAL AND METHODS

### Bacterial strains and growth conditions

*Vibrio cholerae* El Tor C6706 Δ*lacZ*^+^ (Sm^r^), C6706 Δ*lacZ*^−^ (Sm^r^), WT strains, *Vibrio cholerae* C6706 Δ*napA* (Sm^r^), Δ*adhE (SmR)*, Δ*napA adhE* (Sm^r^), Δ*nqrF,* Δ*vc1890* (Sm^r^), Δ*vc1890 nqrF* (Sm^r^), Δ*vc0786* (Sm^r^), Δ*vc0786 nqrF* (Sm^r^), Δ*vc1581*(Sm^r^), Δ*vc1581 nqrF* (Sm^r^), Δ*fdnI* (Sm^r^), Δ*fdnI nqrF* (Sm^r^) and the transposon insertion *napC*, *dsbA*, *tatA*, *moaA*, *narP*, *ccmF* mutant strains, and *Escherichia coli* K12 and the single gene knockout mutant *adhE* from the Keio collection (36) were used in this study. Strains were grown aerobically at 37°C in 15 ml tubes containing 3 ml of LB medium (10 g tryptone, 10 g NaCl, and 5 g yeast extract/L) shaking at 300 rpm. To study NO_3_^−^ reduction-dependent growth arrest and culture viability, 1.5 × 10^7^ aerobically grown cells were inoculated into 15 ml tubes containing 13 ml defined M9 minimal medium with 1% glucose. To study NO_3_^−^ respiration-dependent expansion, complete LB medium buffered with 50 mM phosphate buffer at the indicated pH and supplemented or not with the indicated [KNO_3_] was used. To establish hypoxic conditions, tubes were closed with rubber septa and oxygen removed by sparging the headspace in each tube with argon gas for 3 minutes. In finer-scale analyses for *dsbA* and *nqrF*, we used a plate-reader based growth assay and tracked OD values every 10 minutes using a BioTek Eon plate reader. Culture plates were loaded with the same media and culture concentrations as above and mineral oil was overlaid onto each well along with oxygen-impermeant sealing film.

Where appropriate, antibiotics were added to *V. cholerae* and *E. coli* cultures at the following concentrations: 200 (streptomycin, Sm), 100 (carbenicillin, Cb) and 50 μg/mL (kanamycin, Km).

### Construction of plasmids for V. cholerae mutants

*V. cholerae* mutants were created by allelic exchange with the suicide plasmid pCVD442 (37). To generate Δ*nqrF,* Δ*vc1890, Δvc0786*, Δ*vc1581,* Δ*fdnI* deletion mutants, ca. 1kbp upstream and downstream DNA fragments flanking each coding regions were PCR-amplified with the primers listed in Table S3. Upstream and downstream DNA fragments were spliced together by overlapping PCR. The resulting ca. 2 Kb fragments were digested as shown in table S2 and cloned into pCVD442. Constructs were transformed into *E. coli* DH5α λpir for amplification. They were confirmed by sequencing, transformed into the *E. coli* donor strain SM10 λpir and conjugated for 6 hours at 37°C with *V. cholerae* C6706 by mixing equal volumes (1ml) of exponential phase cultures, and spot-plating. Single crossover *V. cholerae* were selected on LB plates with Sm and Cb. Re-streaked single colonies were then plated on salt-free LB agar containing 10% (w/v) sucrose and Sm. Colonies were streaked on carbenicillin plates to confirm loss of pCVD442 and then checked by PCR for successful deletion mutants.

### Transposon-insertion sequencing (TIS)

TIS was performed essentially as described (Chao et al., 2013) but using oxic and hypoxic *V. cholerae* cultures grown for 6 hours in complete LB medium buffered at pH 8 with 20 mM of KNO_3_ In brief, ~500,000 transposon mutants were generated by conjugation of *V. cholerae* C6706 with SM10λpir *E. coli* carrying the Himar1 suicide transposon vector pSC189 (Chiang SL et al., 2002). After incubation under the conditions explained above, liquid cultures were plated on LB medium agar plates to obtain single isolated colonies, they were collected and their genomic DNA was pooled and analysed on an Illumina MiSeq bench-top sequencer (Illumina, San Diego, CA). Insertion sites (which included 35% of TA sites) were identified as described (Chao et al., 2013), and significance was determined using Con-Artist simulation-based normalization as described (Pritchard et al., 2014). Results were visualized using Artemis (Carver et al., 2012).

### NO_3_^−^ reduction screen of arrayed transposon mutant library

A near-saturating transposon insertion library in *Vibrio cholerae* strain C6706 (Cameron D.E. *et al.,* 2008) consisting of 3,156 insertion mutants was used to identify *V. cholerae* genes required to reduce NO_3_^−^ to NO_2_^−^. C6706 WT and library insertion strains were first grown aerobically in 200 μl of LB in 96-well plates at 37°C for 24 hours. These cultures were then used as inocula into 96-well plates containing 200 μl of buffered (pH 8) complete LB medium, in the presence of 3 mM of NO_3_^−^. The 96-well plates were sealed with aluminium sealing film and incubated without agitation at 37°C for 24 hours. To measure NO_3_ reduction, 96-well plates were centrifuged at 8,000 g for 15 minutes and 100 μl of cell-free supernatant transferred to another 96-well plate. NO_2_^−^ in the supernatant was quantified by the Griess test (8).

### Determination of *in vitro* and *in vivo* nitrate reductase (NR) activity

*V. cholerae* C6706 WT and Δ*nqrF* mutant strains were grown aerobically in LB medium for 24 hours. 1.5 x10^7^ cells from those cultures were then incubated anaerobically in buffered complete LB medium at pH 8 in the presence of 20 mM of NO_3_^−^. These cultures were collected and *in vitro* NR activities were determined as previously described (8). The Griess test was used to determine *in vivo* NR activity by quantifying NO_3_^−^ reduction to NO_2_^−^ in hypoxic culture supernatants. Nitrite concentrations were normalized to the protein levels of each sample by the Bradford assay (Bio-Rad Laboratories).

### Measurement of membrane potential

The flow cytometry-based BacLight Bacterial Membrane Potential Kit (Invitrogen) was used to measure membrane potential. 1mL aliquots from each hypoxic culture were pelleted, washed and resuspended in 600 μl LB medium and then incubated with 30 μM 3,3’-diethyloxacarbocyanine iodide (DiOC2) in the dark for 5 minutes at 20C. 5 μM carbonyl cyanide 3-chlorophenylhydrazone (CCCP) was used as a positive control for membrane depolarization. Fluorescence was measured with a Biorad S3e cell sorter at an excitation wavelength of 488 nm and an emission wavelength of 525/30 nm (green) or 655 nm (red).

### Competition assays

Aerobic overnight cultures of WT *V. cholerae* C6706 *lacZ^+^* and Δ*napA lacZ^−^,* Δ*adhE lacZ^−^* and Δ*napA adhE lacZ^−^* strains were harvested and 1 × 10^7^ cells were co-inoculated at 1:1 ratio in 15 ml tubes containing 13 ml of complete LB medium buffered at pH 8, with or without NO_3_^−^. Tubes were closed with rubber septa and purged of oxygen as described above. After 8 hours of incubation under such conditions at 37°C, an aliquot was diluted and plated on LB agar plates containing X-Gal to enumerate WT and respect the mutant strains present in the culture. Competitive indices (CIs) were determined by dividing the ratio of white (mutant) to blue (WT) colonies by the ratio in the inoculum.

### Streptomycin-treated adult mouse model of *V. cholerae* intestinal colonization

The Sm-treated mouse model was performed identically to our previous report (8). Mice were inoculated with a 1:1 mix of WT and Δ*adhE lacZ^−^* or Δ*napA/adhE lacZ^−^ V. cholerae.* CI values were determined as described above.

### Statistics and reproducibility

Statistical significance was assessed with the Student’s t-test or a Mann-Whitney U test where indicated. A p-value of less than 0.05 was considered statistically significant. Assays were performed with three biological replicates unless otherwise indicated.

## ACKNOWLEDGMENTS

Research in the Cava lab is supported by The Swedish Research Council (VR), The Knut and Alice Wallenberg Foundation (KAW), The Laboratory of Molecular Infection Medicine Sweden (MIMS) and The Kempe Foundation. Research in the Waldor lab is supported by NIH grant RO1AI-042347 and HHMI. EB was supported by a Kempe Foundation fellowship. BS was supported by the Natural Sciences and Engineering Council of Canada (PGSD3-487259-2016). We thank J. J. Mekalanos for the *V. cholerae* C6706 Tn-mutant library.

